# Brain Structural and Resting-state Functional Network Changes Following Expiratory Musculature Targeted Resistance Training in Healthy Young Adults: A Pilot Study

**DOI:** 10.64898/2026.07.09.737407

**Authors:** Rahul Krishnamurthy, Douglas H. Schultz, Yingying Wang, Steven M. Barlow, Angela M. Dietsch

**Affiliations:** Department of Speech and Hearing Sciences, University of New Mexico, Albuquerque, NM, United States; Department of Psychology, University of Nebraska-Lincoln, Lincoln, United States; Center for Brain, Biology, and Behavior, University of Nebraska-Lincoln, Lincoln, United States; Department of Special Education and Communication Disorders, University of Nebraska-Lincoln, Lincoln, United States; Department of Biological Systems Engineering, University of Nebraska-Lincoln, Lincoln, United States

**Keywords:** EMST, sensorimotor plasticity, deglutition, dysphagia

## Abstract

Multimodal imaging approaches that combine structural and functional neuroimaging provide a robust framework for examining neuroplastic adaptations that may not be captured by any single modality. The present study investigated the effects of a four-week expiratory muscle strength training (EMST) program on structural and resting-state functional connectivity in healthy young adults. Five healthy young adult males (aged 19–35 years) completed a standard four-week EMST protocol and underwent pre- and post-training imaging assessments. Structural neuroimaging included T1-weighted and diffusion-weighted MRI, which were analyzed using voxel-based morphometry, surface-based morphometry, and white-matter structural connectivity. Functional neuroimaging consisted of resting-state fMRI to assess training-related changes in functional architecture, network connectivity, and global network measures. Structural MRI analyses revealed no significant changes in gray or white matter volume, cortical morphology, or white-matter structural connectivity following EMST (all FWE- or FDR-corrected *p* > .05). In contrast, resting-state fMRI demonstrated a significant increase in whole-brain functional connectivity (FDR-corrected *p* = .036), accompanied by greater network integration, reflected in increased local efficiency and transitivity and reduced modularity. Network-level analyses showed enhanced within- and between-network connectivity in sensorimotor and cognitive circuits. Our findings demonstrate robust functional reorganization following EMST, despite the absence of detectable macrostructural or large-scale white-matter connectivity changes, at least within the timescale and sample characteristics of the current study. These results reflect early-stage neuroplasticity, both globally and within the networks underlying speech and swallowing control and suggest that functional reorganization occurs early in training and likely precedes longer-term structural modifications in these networks.

## Introduction

Although neuroplasticity is widely acknowledged as the long-term goal of speech-and swallowing-targeted exercises [1,2], two significant gaps have limited progress in this area. First, a lack of quantitative evidence demonstrating neuroplastic changes, and second, a limited understanding of the mechanistic pathways through which targeted exercises may induce such changes. These gaps have limited our ability to precisely design and optimize exercise-based therapies targeting speech and swallowing deficits in aging and disease. In our previous studies, we have demonstrated that speech and swallowing-targeted strength training program, such as the expiratory muscle strength training (EMST) induces functional neuroplasticity through mechanisms similar to those of limb strength training [3–5]. We also provided preliminary quantitative evidence supporting this mechanistic framework, including molecular and functional neuroplastic adaptations following EMST.

Our foundational studies in a small cohort of healthy young adults demonstrated that a four-week EMST program not only increased maximum expiratory pressure (MEP; *p* < .001), as expected, but also elevated circulating serum levels of brain-derived neurotrophic factor (BDNF; *p* = .006) and induced functional neuroplastic changes. Using a swallowing task-based fMRI approach, we observed significantly increased post-training activation (all *p* < .05) in the bilateral primary motor and somatosensory cortices, supplementary motor areas, insula, cerebellum, middle frontal gyrus, right thalamus, and right anterior cingulate cortex [3,4]. While the task-based fMRI provides valuable insights into individual or modular neural activity during task performance, such as swallowing, it does not capture changes in the brain’s intrinsic connectivity architecture or its structural substrate. Because neuroplasticity manifests not only through changes in task-specific neural activity but also through modifications in the brain’s gray and white matter composition and intrinsic (resting-state) communication patterns, the use of complementary neuroimaging modalities is essential.

Structural (T1-weighted) MRI, diffusion-weighted imaging (DW-MRI), and resting-state fMRI (rs-fMRI) together provide a comprehensive multimodal framework for assessing these broader neuroplastic adaptations, enabling characterization of both structural remodeling and intrinsic network reorganization. In this current research note, we aim to investigate the effects of EMST on structural and functional brain networks in healthy young adults using a multimodal neuroimaging approach. The following are our specific research questions.

1. Does EMST induce measurable changes in brain structural features, including gray matter volume, white matter volume, cortical thickness, sulcal depth, fractal dimension and gyrification index in healthy young adults?
2. Does EMST alter white matter structural connectivity and network topology, as assessed using graph-theory metrics at the whole-brain (global) level in healthy young adults?
3. Does EMST modulate resting-state functional connectivity and network topology at the global and network levels, as measured by graph theory metrics within and between major brain networks?
4. In addition to whole-brain analyses, do specific regions of interest (ROIs; nodes) that previously [3,4] showed increased functional activation following EMST (see Table 1) exhibit localized (nodal level) changes in structural and functional connectivity?

**Table 1.**
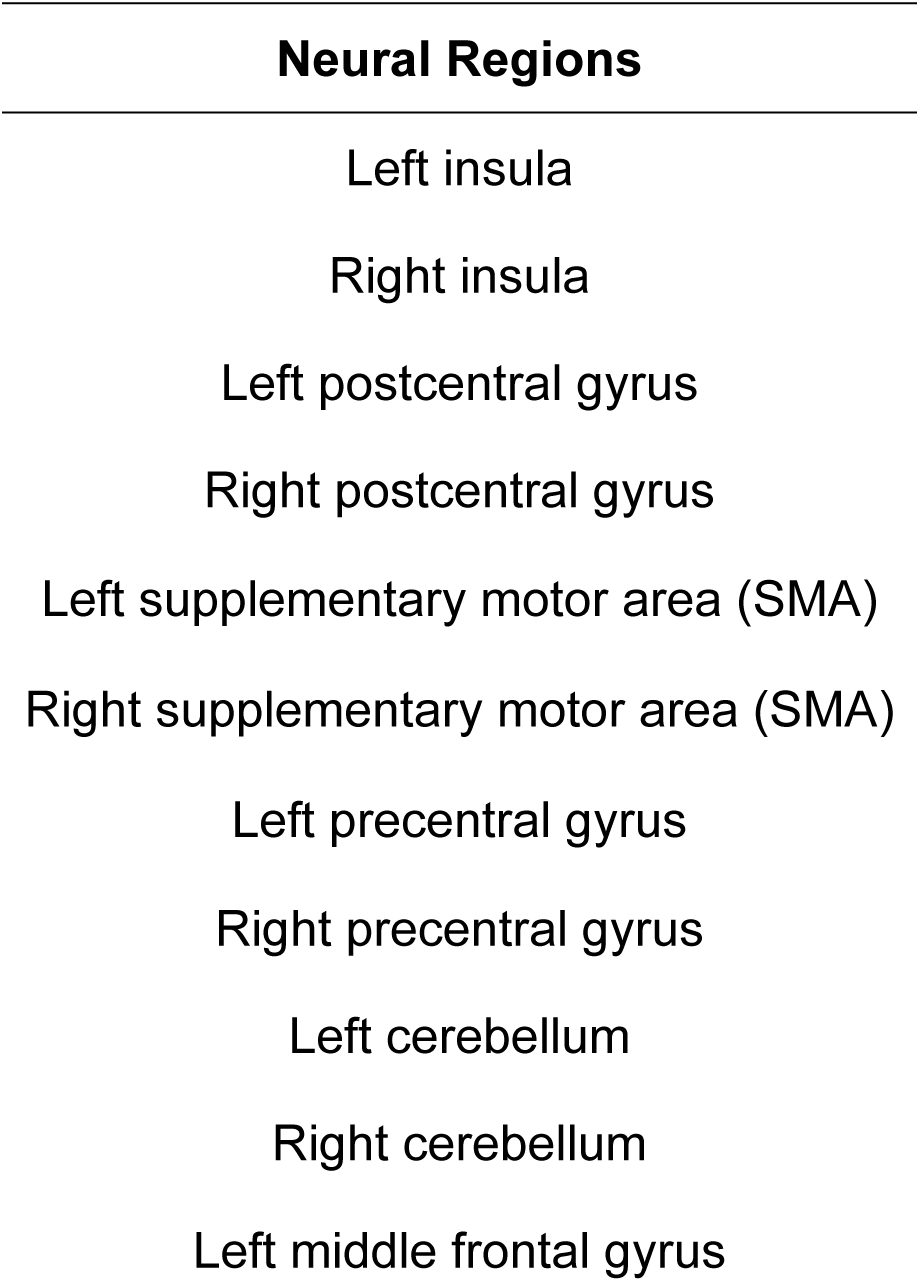
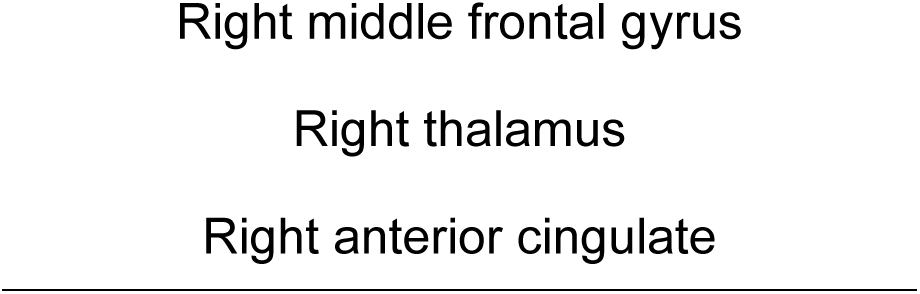
Neural regions of interest previously implicated with EMST [3,4]. *Note*. EMST = Expiratory Muscle Strength Training.

## Methods

### Participants and training

The study protocol was approved by the University of Nebraska–Lincoln institutional review board (#23042), and all participants signed a written informed consent. Five right-handed, healthy young adult men aged 19–35 (Mean = 28.8, S.D. = 2.68) were recruited from the community and participated in this study. All participants had lung volumes and capacities, and body mass index within the normal range. They also had no history of smoking or neurological, psychiatric, pulmonary, speech, or swallowing disorders by self-report. All participants underwent four weeks of expiratory musculature-targeted resistance training following the protocol proposed by Sapienza [6] using the EMST-150 device (Aspire Respiratory Products). Further details of the training specifications (dosage), physiological (MEP), and neuroplastic outcomes have been reported in our previous studies [3,4].

### Multimodal brain imaging parameters and timeline

For each participant, high-resolution T1-weighted anatomical, rs-fMRI, DW-MRI and field map scans were acquired on a Siemens 3T Magnetom Skyra. T1-weighted structural images were obtained using a magnetization-prepared rapid acquisition gradient echo (MPRAGE) sequence with the following parameters: TI/TR/TE = 991ms/2.2s s/3.37ms, flip angle = 7°, voxel size = 1 mm³, bandwidth = 200 Hz/pixel, and GRAPPA acceleration factor of 2 in the anterior-posterior direction. DW-MRI data were acquired using a Spin-Echo EPI sequence with parameters: flip angle/TE/TR/TA/FOV = 90 degrees/88 ms/8320 ms/5:59 min/256 × 256 mm; with b = 1000 s/mm² in 60 diffusion gradient directions. Each rs-fMRI data set was acquired using a multiband whole-brain echo planar imaging (EPI) sequence (TR = 720 ms, TE = 33.1 ms, flip angle = 52°, multiband factor = 8, voxel size = 2.0 mm isotropic, 72 slices) for 16 minutes while participants rested with eyes open, fixating on a crosshair. The baseline scans were obtained one day before the beginning of the training, and the post-training scans were obtained within three days of completing the training.

### Multimodal brain imaging pre-processing and analysis pipelines

All preprocessing and analyses were conducted on the University of Nebraska–Lincoln’s SWAN supercomputer cluster. Before any preprocessing steps, all structural T1-weighted and DW-MRI images were visually inspected to ensure data quality, and no scans were excluded due to significant artifacts or abnormalities. We converted the multimodal (structural and resting-state functional) DICOMs to NIFTI format using the dcm2niix tool (https://github.com/rordenlab/dcm2niix). **Fig 1** shows the multimodal analyses pipeline, and the details of modality-specific pre-processing and analyses are described in the sections below.

**Fig 1.**
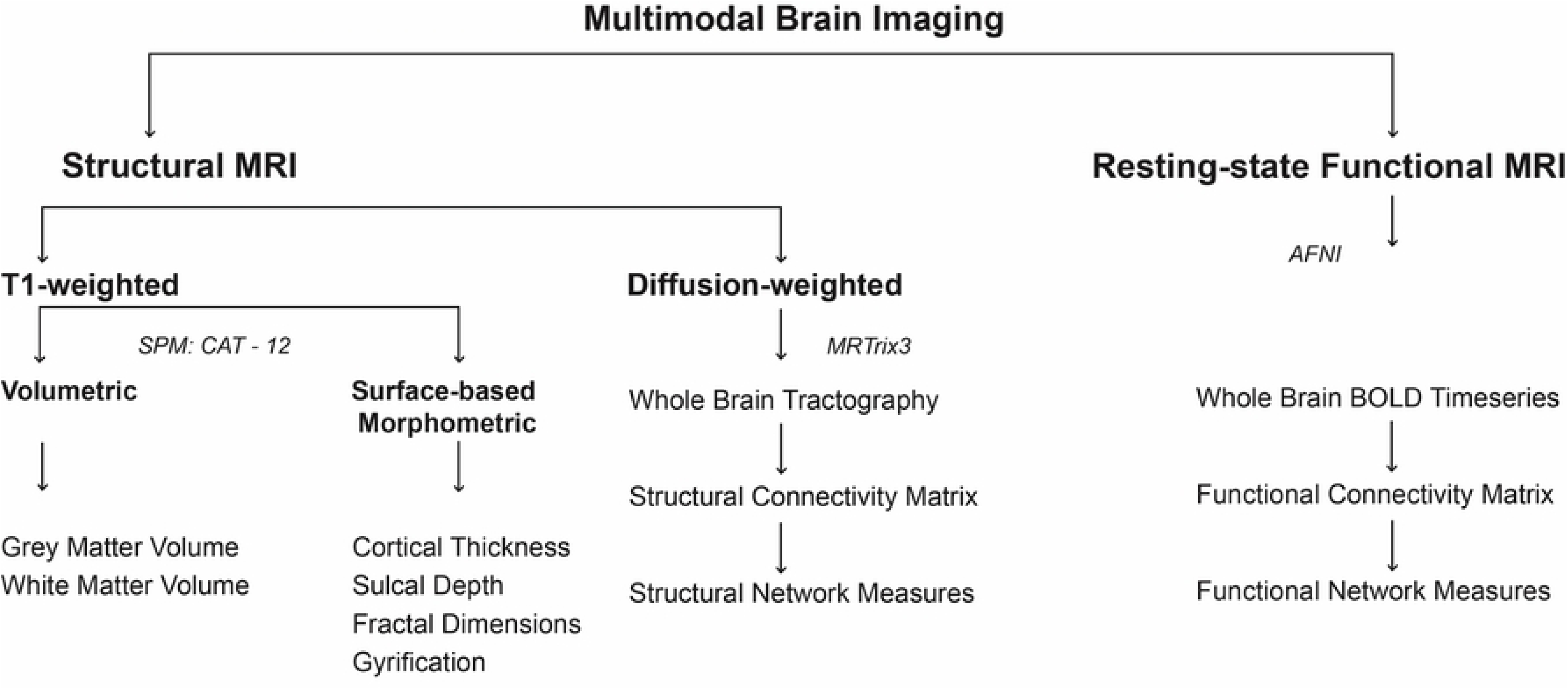
Multimodal neuroimaging pre-processing and analyses pipelines used in the current study.

#### T1-weighted volumetric and surface-based morphometric data

The T1-MPRAGE structural images were pre-processed using the Computational Anatomy Toolbox (CAT12; http://www.neuro.uni-jena.de/cat/) [7] implemented in SPM12 (Wellcome Centre for Human Neuroimaging, London, UK). Furthermore, these images underwent anatomical reconstruction and segmentation using the Freesurfer pipeline [8,9] to obtain the Desikan-Killiany parcellation atlas [8] for each subject (file: aparc+aseg.mgz). These Freesurfer parcellations were also used to generate the five-tissue-type (5tt) segmented image required to perform anatomically constrained tractography [9] using the *5ttgen* function in MRtrix3 [10].

We followed the CAT12 standard protocol [7] for preprocessing voxel-based and surface-based morphometric data. This preprocessing pipeline included bias correction, segmentation of gray matter, white matter, and cerebrospinal fluid, spatial normalization to MNI space, and modulation. For voxel-based morphometry, the modulated gray matter and white matter volumetric maps were smoothed with an 8 mm full-width at half maximum (FWHM) Gaussian kernel. For surface-based morphometry, we extracted additional surface parameters, including cortical thickness [12], sulcal depth, gyrification index [13], and fractal dimensions [14]. Cortical thickness data were smoothed with a 15 mm FWHM, and sulcal depth, gyrification index, and fractal dimension data were smoothed with a 20 mm FWHM of the Gaussian kernel [7]. We further extracted total intracranial volumes for each participant, and no scans were excluded due to poor quality. The smoothed data were used to perform paired sample t-tests for statistical comparisons between baseline and post-training conditions for gray matter and white matter volume, cortical thickness, sulcal depth, gyrification index, and fractal dimension in the CAT12 and SPM12 statistical modules.

Each participant’s total intracranial volume was used as a covariate for voxel-based morphometry analysis to correct for different brain sizes, and their age and sex were included for both voxel-based morphometry and surface-based morphometric analyses. A threshold of *p* < .05 was applied to all analyses after family-wise error (FWE) correction for multiple comparisons (cluster-level) to compare baseline and post-training conditions. Only those clusters with > 20 voxels and regional overlap exceeding 40% for gray matter and white matter volumetric data were retained. After the statistical analysis, we exported the voxel-based and surface-based morphometric results from CAT12 and visualized them in SPM12. Regional brain volumetric differences were identified using the neuromorphometric atlas, and the surface-based region-specific differences were labeled using the Desikan-Killiany atlas [8] within CAT12’s results viewer.

#### DW-MRI pre-processing

The DW-MRI was pre-processed and analyzed using a custom pipeline based on DIPY (1.11.0) [15], FMRIB Software Library (FSL 6.0.7.17) [16], and MRtrix3 [11]. Pre-processing steps included (1) quality control check to remove DW-MRI volumes with artifacts (*DTIPrep 1.2.11;* https://www.nitrc.org/projects/dtiprep), (2) denoising using overcomplete local principal component analysis (*DIPY*), (3) removal of intensity oscillations (Gibbs unringing) (*DIPY*), (4) bias field corrections (*ANTs*), (5) between-volume motion correction (*DIPY*), and (6) eddy current corrections using *eddy* (*FSL*). Further details of DW-MRI pre-processing are described in our previous works [3,17]. This pre-processed DW-MRI data was used to perform anatomically constrained tractography, generate the structural connectome and obtain network-based structural connectivity measures.

#### Tractography

We performed anatomically constrained tractography [10] and obtained structural connectomes using the pipeline described by Tahedl et al. [18] implemented within MRtrix3. The pre-processed DW-MRI data were used for brain mask generation and non-brain tissue removal using *dwi2mask* (MRtrix3). Next, the five-tissue-type (5tt) segmented image, generated in a prior step, was co-registered to the DW-MRI space using non-linear co-registration via ANTs. We then estimated the response function using *dwi2response* with the *msmt_5tt* algorithm, and fiber orientation distributions (FODs) were computed using the *dwi2fod* function with the *msmt_csd* algorithm. T1-weighted MRl and 5tt images were also co-registered using a rigid + affine transformation algorithm in ANTs. The *tckgen* (MRtrix3) algorithm with a second-order integration over FOD (using the *iFOD2* algorithm) was used for fiber tracking and tractography.

We generated 10 million streamlines with the following parameters: minimum length = 4 mm, maximum length = 200 mm, maximum angle between successive steps = 45°, unidirectional tracking with the backtracking option enabled. Streamlines were seeded from the gray matter/white matter interface using the previously generated 5tt images. Finally, the structural connectome was constructed from the FreeSurfer atlas using the *tck2connectome* (MRtrix3) function. The *tcksift2* algorithm (MRtrix3) was also used to optimize streamlines and fixel-wise fiber densities to obtain a whole-brain tractogram. Finally, the freesurfer parcellations (84-region Desikan-Killiany atlas) were converted into MRtrix3-compatible format using *labelconvert,* and whole-brain structural connectivity matrices were generated for each subject using *tck2connectome*. The resulting matrices represent weighted, undirected graphs, where nodes correspond to anatomical regions and edge weights represent the SIFT2-weighted connection strength (i.e., the fiber density–weighted sum of streamlines) between each pair of regions. Finally, we computed the mean structural connectivity for the whole brain and within predefined ROIs (Table 1).

#### Resting-state fMRI pre-processing

The rs-fMRI data were preprocessed using the Analysis of Functional Neuroimages (AFNI)[19] and the FreeSurfer pipelines [8,9]. The AFNI preprocessing pipeline was implemented using the *afni_proc.py* script, which involved a series of processing blocks, each with specific parameters tailored to optimize data quality and minimize artifacts. First, the raw rs-fMRI data underwent despiking to remove large amplitude spikes. Slice timing correction was then performed to account for differences in slice acquisition times. After that, the functional data were aligned to a common space (MNI152_2009c) using affine and non-linear transformations.

Motion artifacts were corrected through volume registration, aligning each volume to a reference volume. AFNI’s default minimum-outlier volume was used as the reference, and motion parameters were estimated and applied during alignment. Motion scrubbing was implemented to remove time points with excessive motion using a threshold of 0.3 mm to improve the data quality further. Additionally, spatial smoothing using a 4 FWHM Gaussian kernel was applied to improve the signal-to-noise ratio, and intensity normalization was used to ensure consistency in signal intensity across participants. A brain mask was created for each participant based on their Freesurfer parcellated anatomical data to focus the analysis solely on brain voxels. Moreover, to reduce the impact of physiological noise, nuisance signals from white matter, ventricles, and the whole brain were regressed from the functional data. A bandpass filter (0.01 - 0.1 Hz) was applied to remove high and low frequency noise. Finally, we extracted the mean BOLD signal time series at a whole-brain level using the 360 cortical regions of the Glasser multimodal parcellation atlas [20]. Pearson correlations were calculated between all pairs of time series, and Fisher’s z-transformation was then applied to construct a functional connectivity matrix.

Mean functional connectivity was computed separately for whole-brain and ROI-based analyses (Table 1). For the whole-brain analysis, we used the 360-region Glasser atlas and functional connectivity values were calculated between all Glasser parcels and averaged to yield a single whole-brain mean connectivity value per participant. These values were then averaged across participants for group-level analyses. Analyses involving predefined ROIs were conducted using the same Glasser atlas as the whole-brain analysis. Predefined ROIs were operationalized as specific Glasser parcels corresponding to regions reported in prior work (shown in Table 1). These parcels were selected from the 360-region Glasser atlas and analyzed separately for targeted connectivity analyses. Thus, whole-brain connectivity was computed across all Glasser parcels, whereas ROI-based analyses were restricted to a subset of Glasser parcels representing the predefined regions of interest.

### Graph theory metrics

Individual structural connectivity matrices derived from the MRtrix3 pipeline and functional connectivity matrices derived from the AFNI pipeline were used to obtain network-based graph-theoretic metrics using the Brain Connectivity Toolbox [21] and NetworkX in Python. Because global signal regression yields a substantial proportion of negative correlations, we retained these negative FC values and used signed weighted matrices for graph construction, preserving the complete FC structure. Edges with zero weight were removed for both modalities, and undirected weighted (signed for FC) graphs were constructed for computing global network metrics. We obtained density, transitivity, average clustering coefficient, global and local efficiency, assortativity, and modularity. Nodal-level metrics included nodal strength, clustering, and betweenness. All metrics were computed per subject and saved for subsequent group-level statistical analysis.

### Resting-state Functional Network Mapping and Network-level Analyses

We mapped the 360 Glasser regions to the seven-network canonical functional parcellation defined by Yeo et al. [22] to analyze resting-state functional connectivity changes at the network level. These networks include the sensorimotor, frontoparietal, dorsal attention, ventral attention (salience), visual, default mode, and limbic networks. For each subject, the Glasser-level functional connectivity matrix was transformed into a 7 × 7 network-level functional connectivity matrix by averaging all Fisher z-transformed correlations between pairs of regions belonging to each pair of networks. Furthermore, we compared mean functional connectivity within and between these networks to investigate how EMST modulated brain networks.

### Statistical analyses

Paired-samples t-tests were used to compare baseline and post-training conditions to investigate changes in brain volumetric and surface-based morphometric features, white matter structural connectivity, functional connectivity, and graph-theory network measures. A family-wise error (FWE) or a false discovery rate (FDR) correction method was applied to control for multiple comparisons.

## Results

### T1-weighted volumetric changes

Voxel-based morphometric analyses showed no significant differences between baseline and post-training conditions in gray matter volume or white matter volume at the whole-brain level (FWE-corrected; *p* < .05). Although gray matter and white matter volumetric measures showed a small decrease post-training, these changes did not reach statistical significance. Table 2 presents the descriptive statistics of total gray matter, white matter volume, and intracranial volume for baseline and post-training conditions.

**Table 2.** Mean (standard deviation) of voxel-based morphometric measures across baseline and post-training conditions.

| Volume (mm<sup>3</sup>) | Baseline | Post-training | %Change |
| --- | --- | --- | --- |
| GMV | 734.6 (28.43) | 726.4 (25.75) | -1.11 |
| WMV | 566.9 (48.33) | 565 (46.29) | -.33 |
| TIV | 1566 (100.7) | 1567 (98.42) | .06 |

*Note.* Voxel-based morphometric measures are reported as absolute volumes. GMV = gray matter volume; WMV = white matter volume; TIV = total intracranial volume.

In addition to whole-brain analyses, we examined regional gray matter and white matter volumetric changes within atlas-defined neural regions previously shown to exhibit increased functional activation following EMST. These regions were defined using CAT12 native atlases (neuromorphometric and Desikan–Killiany). This ROI-based analysis also revealed no significant differences between baseline and post-training conditions. Table 3 presents descriptive statistics and results of paired-sample t-tests comparing volumetric measures across these regions of interest.

**Table 3.**
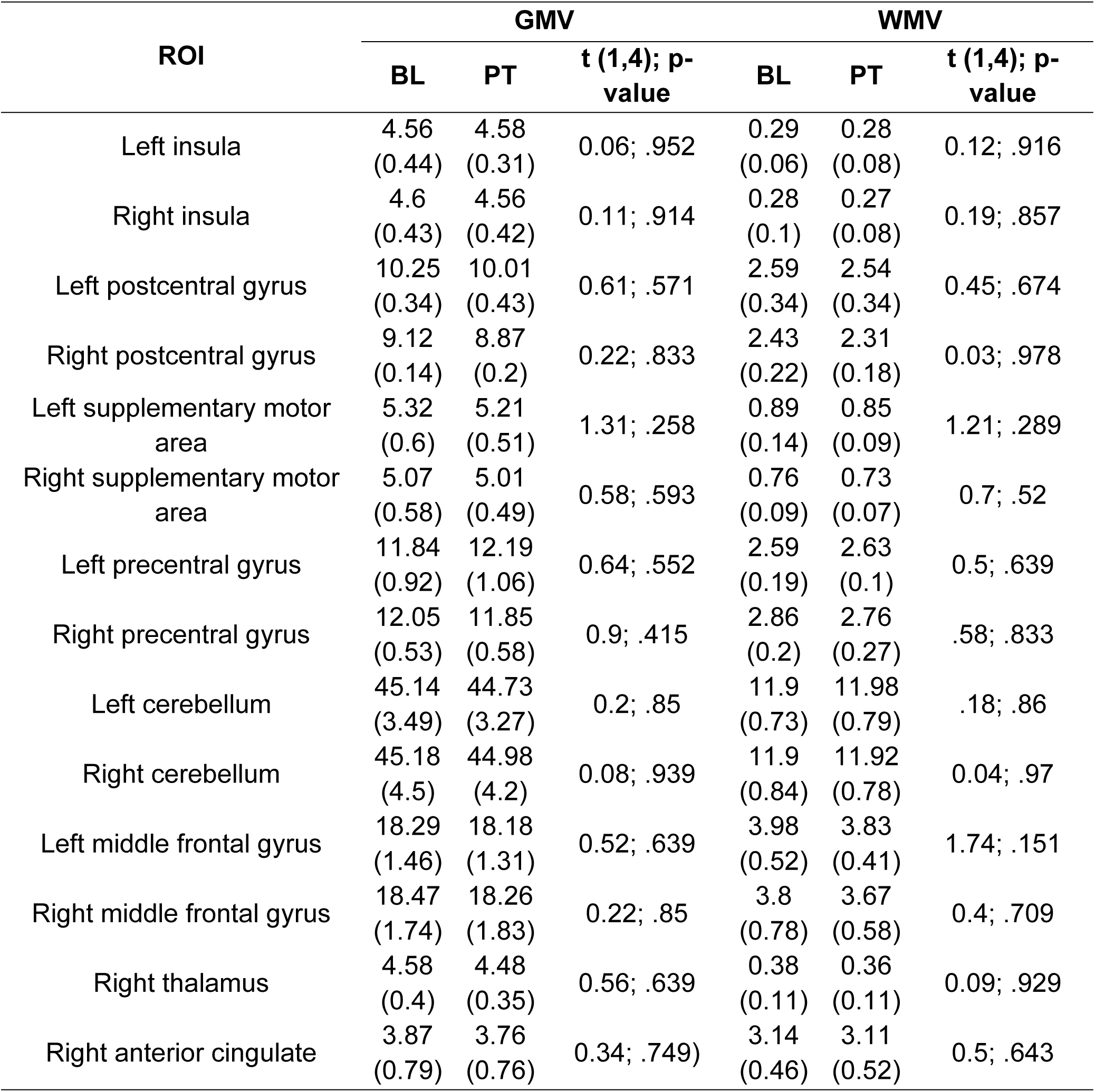
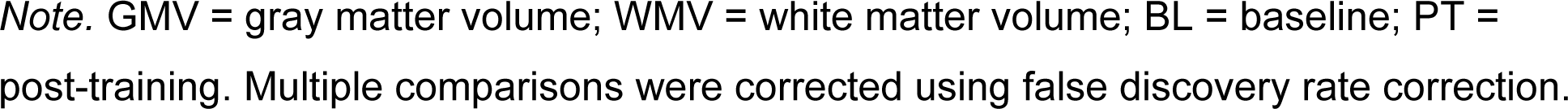
Mean, standard deviation and results of the paired t-test comparing gray matter and white matter volumes for the study’s region of interest (ROIs) across baseline and post-training conditions.

| ROI | GMV |  |  | WMV |  |  |
| --- | --- | --- | --- | --- | --- | --- |
|  | BL | PT | t (1,4); p-value | BL | PT | t (1,4); p-value |
| Left insula | 4.56<br>(0.44) | 4.58<br>(0.31) | 0.06; .952 | 0.29<br>(0.06) | 0.28<br>(0.08) | 0.12; .916 |
| Right insula | 4.6<br>(0.43) | 4.56<br>(0.42) | 0.11; .914 | 0.28<br>(0.1) | 0.27<br>(0.08) | 0.19; .857 |
| Left postcentral gyrus | 10.25<br>(0.34) | 10.01<br>(0.43) | 0.61; .571 | 2.59<br>(0.34) | 2.54<br>(0.34) | 0.45; .674 |
| Right postcentral gyrus | 9.12<br>(0.14) | 8.87<br>(0.2) | 0.22; .833 | 2.43<br>(0.22) | 2.31<br>(0.18) | 0.03; .978 |
| Left supplementary motor area | 5.32<br>(0.6) | 5.21<br>(0.51) | 1.31; .258 | 0.89<br>(0.14) | 0.85<br>(0.09) | 1.21; .289 |
| Right supplementary motor area | 5.07<br>(0.58) | 5.01<br>(0.49) | 0.58; .593 | 0.76<br>(0.09) | 0.73<br>(0.07) | 0.7; .52 |
| Left precentral gyrus | 11.84<br>(0.92) | 12.19<br>(1.06) | 0.64; .552 | 2.59<br>(0.19) | 2.63<br>(0.1) | 0.5; .639 |
| Right precentral gyrus | 12.05<br>(0.53) | 11.85<br>(0.58) | 0.9; .415 | 2.86<br>(0.2) | 2.76<br>(0.27) | .58; .833 |
| Left cerebellum | 45.14<br>(3.49) | 44.73<br>(3.27) | 0.2; .85 | 11.9<br>(0.73) | 11.98<br>(0.79) | .18; .86 |
| Right cerebellum | 45.18<br>(4.5) | 44.98<br>(4.2) | 0.08; .939 | 11.9<br>(0.84) | 11.92<br>(0.78) | 0.04; .97 |
| Left middle frontal gyrus | 18.29<br>(1.46) | 18.18<br>(1.31) | 0.52; .639 | 3.98<br>(0.52) | 3.83<br>(0.41) | 1.74; .151 |
| Right middle frontal gyrus | 18.47<br>(1.74) | 18.26<br>(1.83) | 0.22; .85 | 3.8<br>(0.78) | 3.67<br>(0.58) | 0.4; .709 |
| Right thalamus | 4.58<br>(0.4) | 4.48<br>(0.35) | 0.56; .639 | 0.38<br>(0.11) | 0.36<br>(0.11) | 0.09; .929 |
| Right anterior cingulate | 3.87<br>(0.79) | 3.76<br>(0.76) | 0.34; .749 | 3.14<br>(0.46) | 3.11<br>(0.52) | 0.5; .643 |

*Note.* GMV = gray matter volume; WMV = white matter volume; BL = baseline; PT = post-training. Multiple comparisons were corrected using false discovery rate correction.

### T1-weighted surface-based morphometric changes

Surface-based morphometric analyses revealed no significant differences between baseline and post-training conditions in cortical thickness, sulcal depth, gyrification, or fractal dimensions at the whole-brain level (all FWE-corrected; *p* < .05). Additionally, an ROI-based analysis also showed no significant differences between the two conditions. S1 Table presents the means and standard deviations of surface-based morphometric measures for these ROIs.

### White matter structural connectivity changes

The results of our paired t-test comparisons indicated no significant differences in mean structural connectomes across conditions for either whole-brain [t(1,4) = 0.961; FDR corrected *p* = .39] or predefined ROIs [t(1,4) = 1.11; FDR corrected *p* = .32]. **Fig 2(A)** shows the averaged whole-brain structural connectomes at baseline and post-training, while **Fig 2(B)** shows the structural connectomes for the predefined ROIs, along with bar graphs comparing mean structural connectivity across conditions.

**Fig 2.**
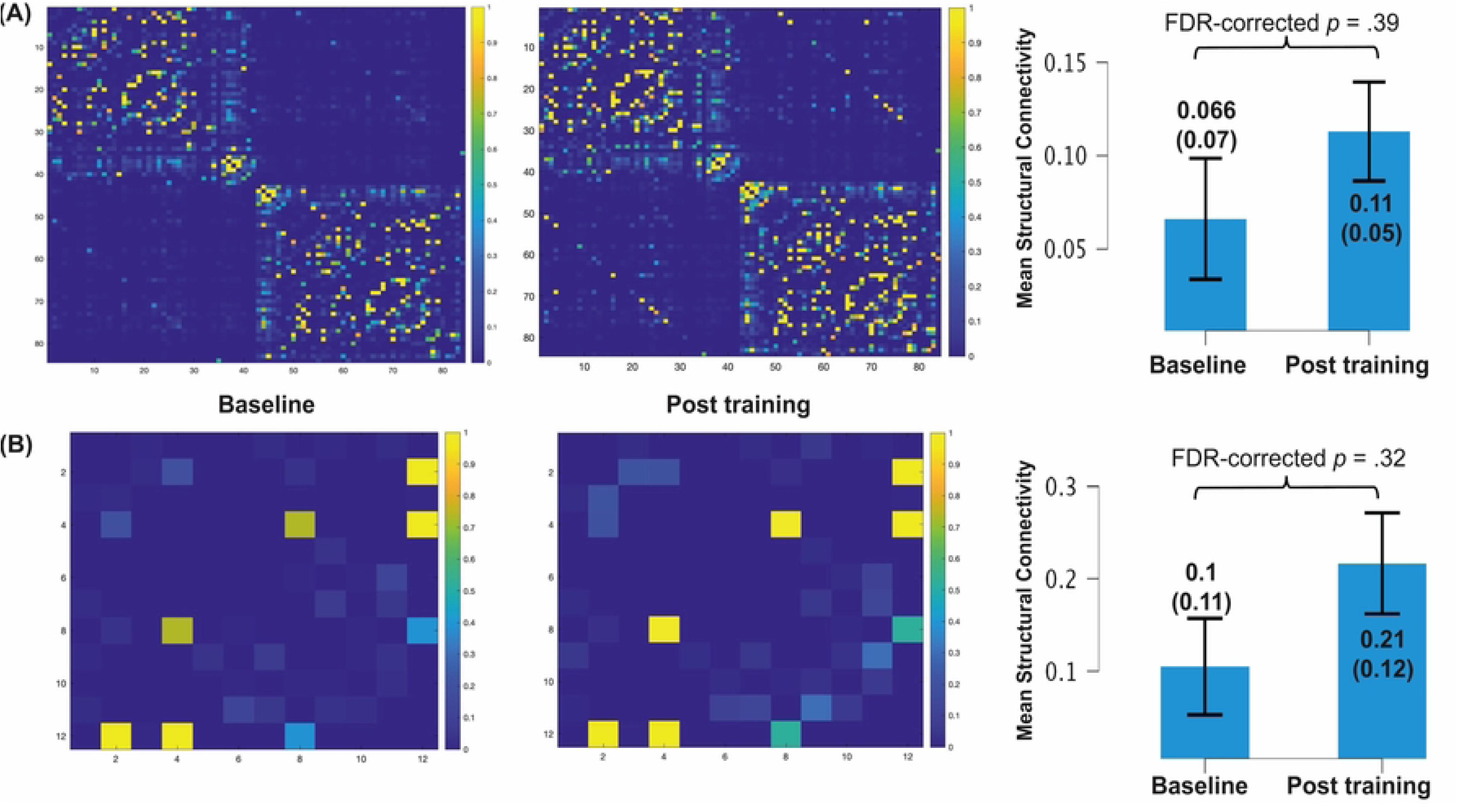
Averaged whole-brain (A) and (B) ROI-based structural connectomes at baseline and post-training. *Note.* Edge weights represent fiber density–weighted streamline strength (SIFT2-weighted). Bar graphs show mean (S.D.) structural connectivity across conditions, and whiskers represent standard error.

Furthermore, when we compared graph-theory global and nodal metrics, none of the network-level measures differed significantly between conditions (FDR-corrected all *p* > .05). Table 4 presents the descriptive statistics and results of paired-sample t-tests for these global metrics, and **Fig 3** illustrates these comparisons using box plots for each measure across conditions.

**Fig 3.**
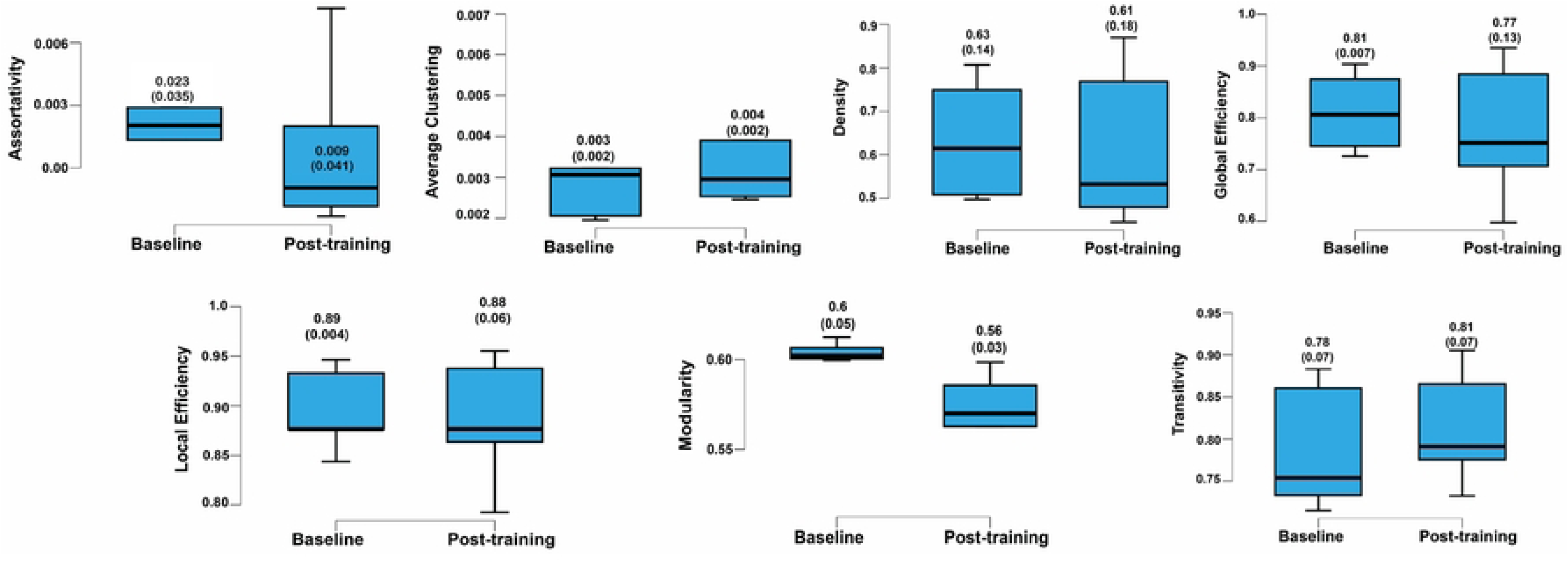
Changes in structural connectome-derived global graph-theory metrics across baseline and post-training conditions.

**Table 4.**
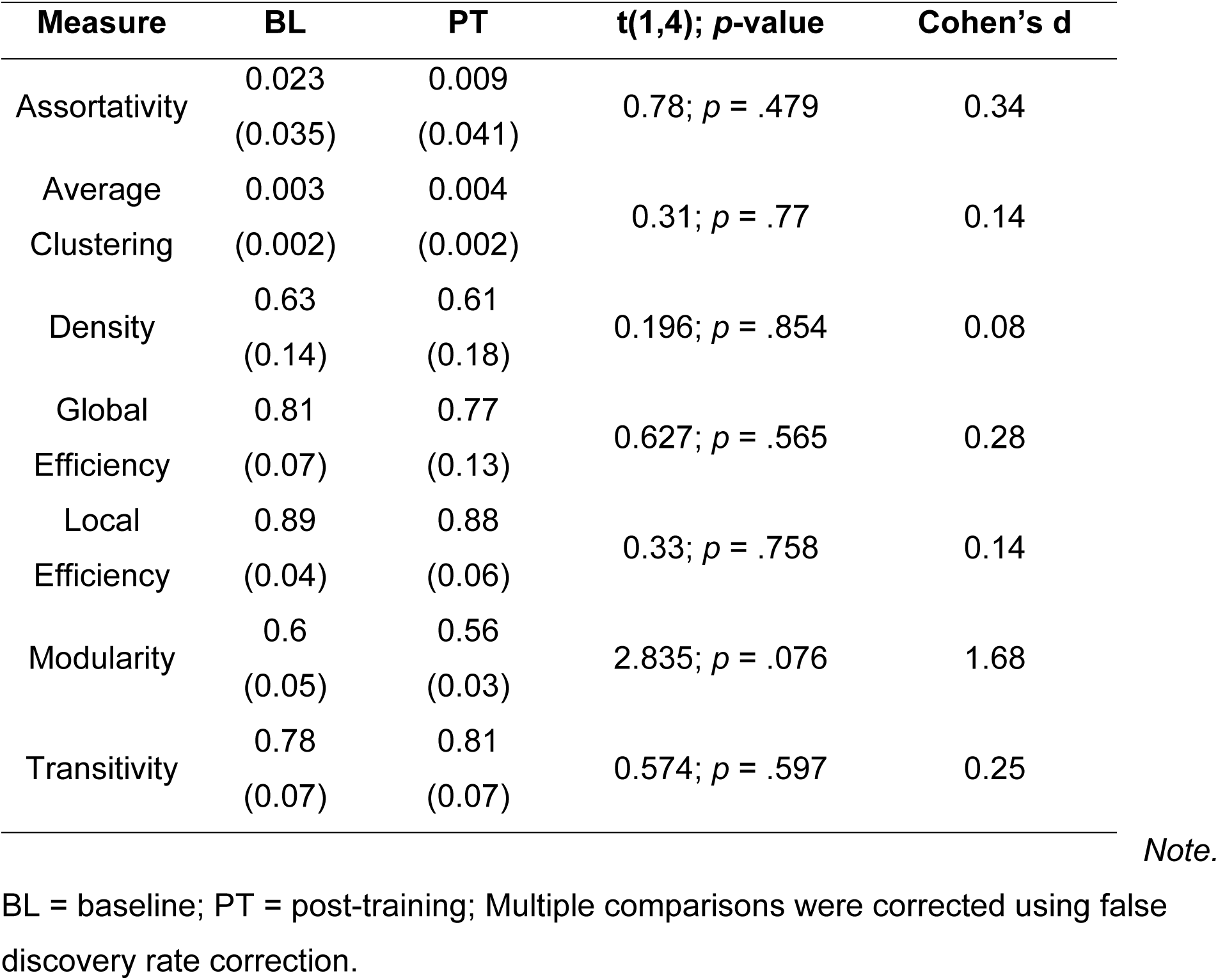
Mean, standard deviation and results of the paired t-test comparing global structural connectivity network measures across baseline and post-training conditions.

S2 Table presents the descriptive statistics and the paired t-test results for each nodal measure. Together, these findings indicate that both whole-brain (global) and region-specific structural connectivity remained stable from baseline to post-training, suggesting no measurable changes in structural connectome organization due to EMST within our small group and study time frame.

### Resting-state functional connectivity changes

The results of our paired t-test comparisons revealed a significant increase in whole-brain mean resting-state functional connectivity from baseline to post-training [t(1,4) = 2.55, FDR-corrected *p* = .036]. Whole-brain mean connectivity was defined as the average of all pairwise functional connectivity values across the 360 × 360 Glasser-based connectivity matrices for each participant. However, no significant difference in the mean functional connectome was observed for predefined ROIs [t(1,4) = 1.11; FDR-corrected *p* = .402]. **Fig 4(A)** shows the averaged whole-brain functional connectomes at baseline and post-training, while **Fig 4(B)** shows the functional connectomes for the predefined ROIs, along with bar graphs comparing mean structural connectivity across conditions.

**Figure 4.**
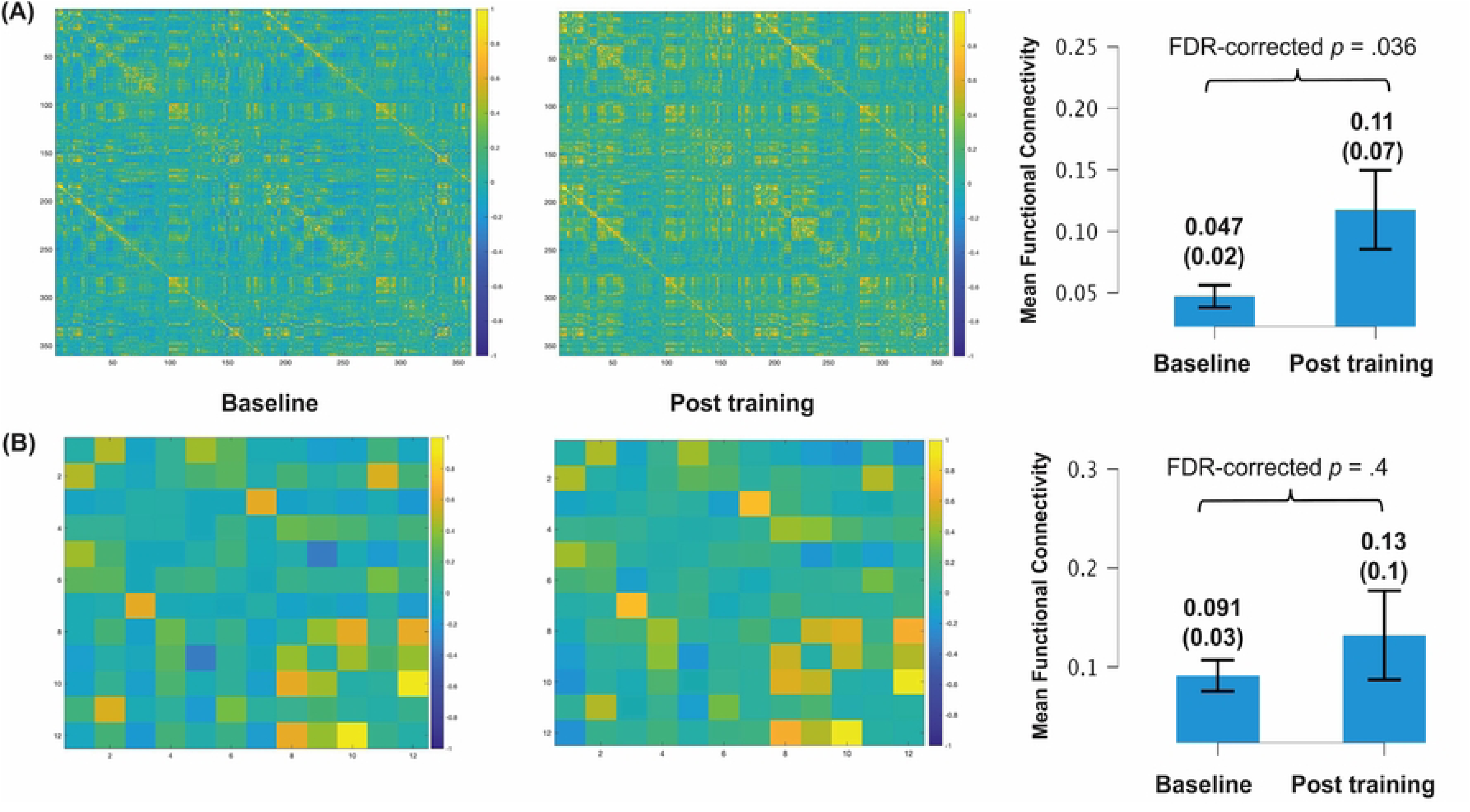
(A) Averaged whole-brain resting-state functional connectomes at baseline and post-training. *Note.* Edge weights represent Fisher’s Z-transformed correlation coefficients between regional BOLD time series. (B) Bar graphs show mean (S.D.) functional connectivity across conditions, and whiskers represent standard error.

Furthermore, when we compared global (whole-brain) graph-theory metrics, we observed trends toward increased network integration following EMST. Specifically, local efficiency (FDR-corrected *p* = .046) and transitivity (FDR-corrected *p* = .031) increased, while modularity decreased significantly (FDR-corrected *p* = .014). In addition, average clustering, density, assortativity, and global efficiency showed increased trends at post-training compared to baseline but did not reach statistical significance (all FDR-corrected *p* > .05). Table 5 shows the descriptive statistics and results of paired-sample t-tests for these global metrics, and **Fig 5** illustrates the comparisons using box plots for each measure at both conditions.

**Fig 5.**
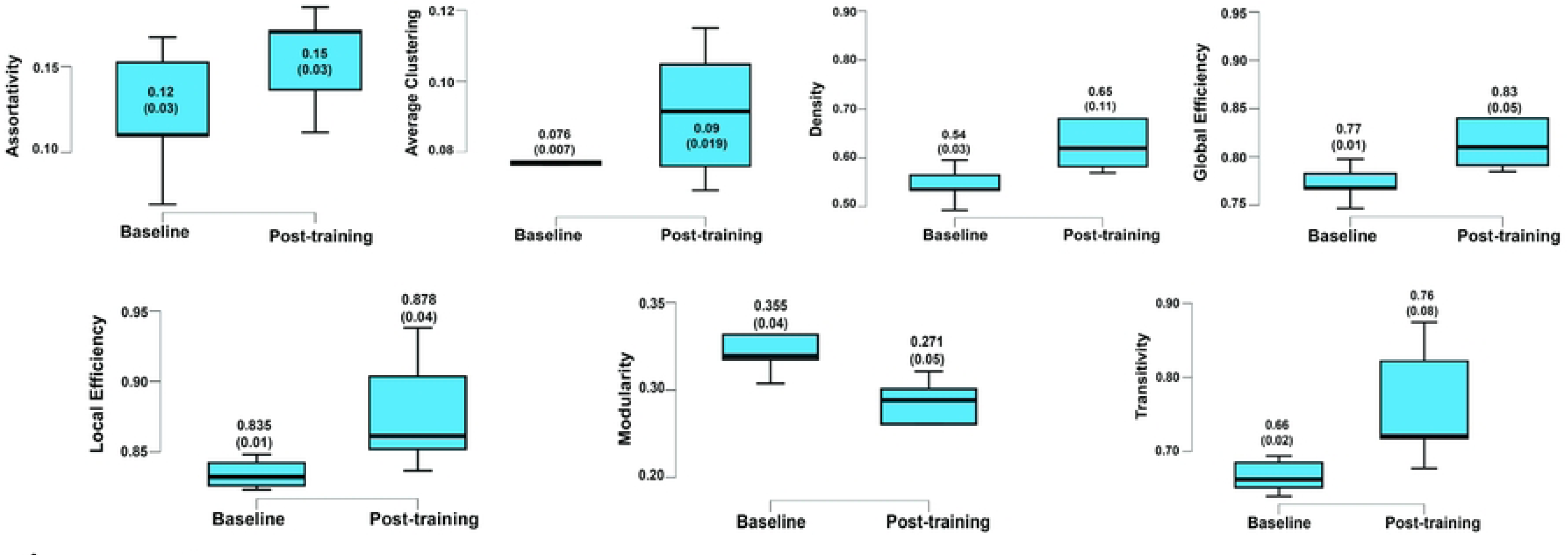
Changes in resting-state functional connectome-derived global graph-theory metrics across baseline and post-training conditions.

**Table 5.** Mean, standard deviation and results of the paired t-test comparing global resting-state functional connectivity network measures across baseline and post-training conditions.

| Measure | BL | PT | t(1,4); *p*-value | Cohen's *d* |  |
| --- | --- | --- | --- | --- | --- |
| Assortativity | 0.12<br>(0.03) | 0.15<br>(0.03) | 1.818; *p* = .143 | 0.813 |  |
| Average<br>Clustering | 0.076<br>(0.007) | 0.09<br>(0.019) | 2.368; *p* = .07 | 1.059 |  |
| Density | 0.54<br>(0.03) | 0.65<br>(0.11) | 2.597; *p* = .06 | 1.161 |  |
| Global<br>Efficiency | 0.77<br>(0.01) | 0.83<br>(0.05) | 2.59; *p* = .061 | 1.16 |  |
| Local<br>Efficiency | 0.835<br>(0.01) | 0.878<br>(0.04) | 2.852; *p* = .046 | 1.276 |  |
| Modularity | 0.355<br>(0.04) | 0.271<br>(0.05) | 4.143; *p* = .014 | 1.853 | **Note.** |
| Transitivity | 0.66<br>(0.02) | 0.76<br>(0.08) | 3.253; *p* = .031 | 1.455 | BL = |

baseline; PT = post-training; Multiple comparisons were corrected using false discovery rate correction.

We examined nodal (regional) graph-theory metrics to determine whether EMST was associated with localized changes in network topology across predefined ROIs. Across regions, nodal strength, clustering coefficient, and betweenness centrality generally showed higher mean values at post-training compared to baseline, suggesting trends toward increased regional connectivity and integration following EMST. However, none of the regional effects survived FDR-correction for multiple comparisons (all FDR-corrected *p* > .05). At the uncorrected level, several regions demonstrated trend-level or nominally significant increases in nodal strength and clustering, including the left precentral gyrus and right anterior cingulate, though these effects did not remain significant after correction. Betweenness centrality showed no consistent pattern of change across regions. Descriptive statistics and paired-sample t-test results for all nodal metrics are presented in S3 Table.

At the within-network level, EMST significantly increased resting-state functional connectivity in the sensorimotor [t(1,4) = 4.562; FDR-corrected *p* < .001], frontoparietal [t(1,4) = 2.961; FDR-corrected *p* = .004], and dorsal attention [t(1,4) = 2.409; FDR-corrected *p* = .003] networks. **Fig 6** shows their box plots at baseline and post-training conditions.

**Fig 6.**
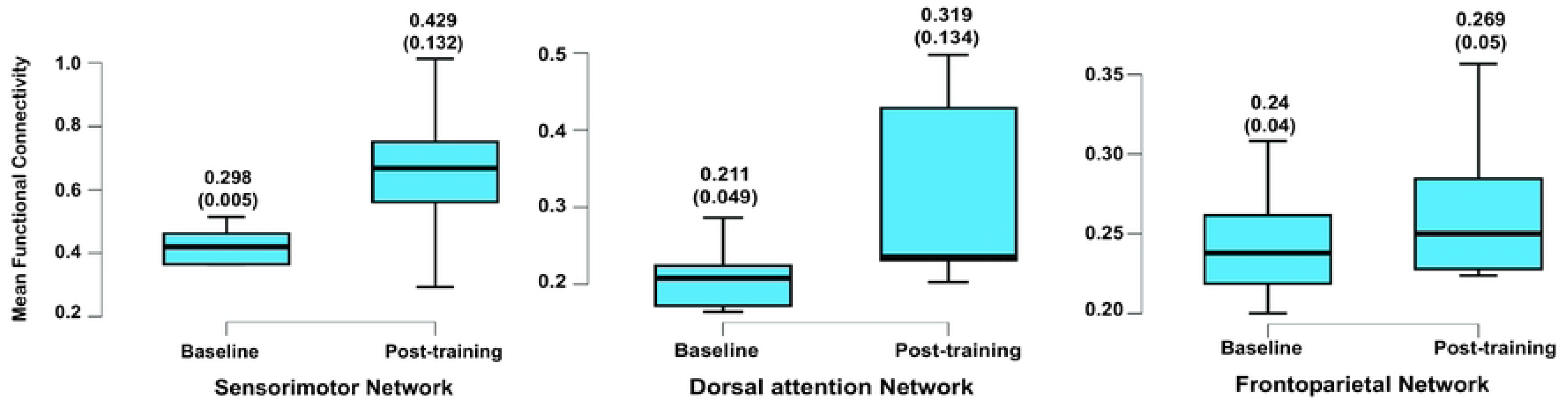
Changes in within-network mean resting-state functional connectivity across baseline and post-training conditions.

At the between-network level, significant post-training increases were observed between the following pairs: sensorimotor and dorsal attention network [t(1,4) = 4.572; FDR-corrected *p* < .001], sensorimotor and frontoparietal network [t(1,4) = 4.79; FDR-corrected *p* = .008], sensorimotor and default mode [t(1,4) = 2.967; FDR-corrected *p* = .004], dorsal attention and ventral attention (salience) networks [t(1,4) = 4.839; FDR-corrected *p* < .008]. Additional significant increases were observed among visual and dorsal attention [t(1,4) = 3.04; FDR-corrected *p* = .003], and visual and frontoparietal networks [t(1,4) = 2.96; FDR-corrected *p* = .004]. **Fig 7** shows the string plot for these increased connections. Overall, these findings indicate that EMST strengthens both local (within-network) and distributed (between-network) connectivity, supporting increased functional integration across sensorimotor, attentional and higher-order cognitive networks.

**Fig 7.**
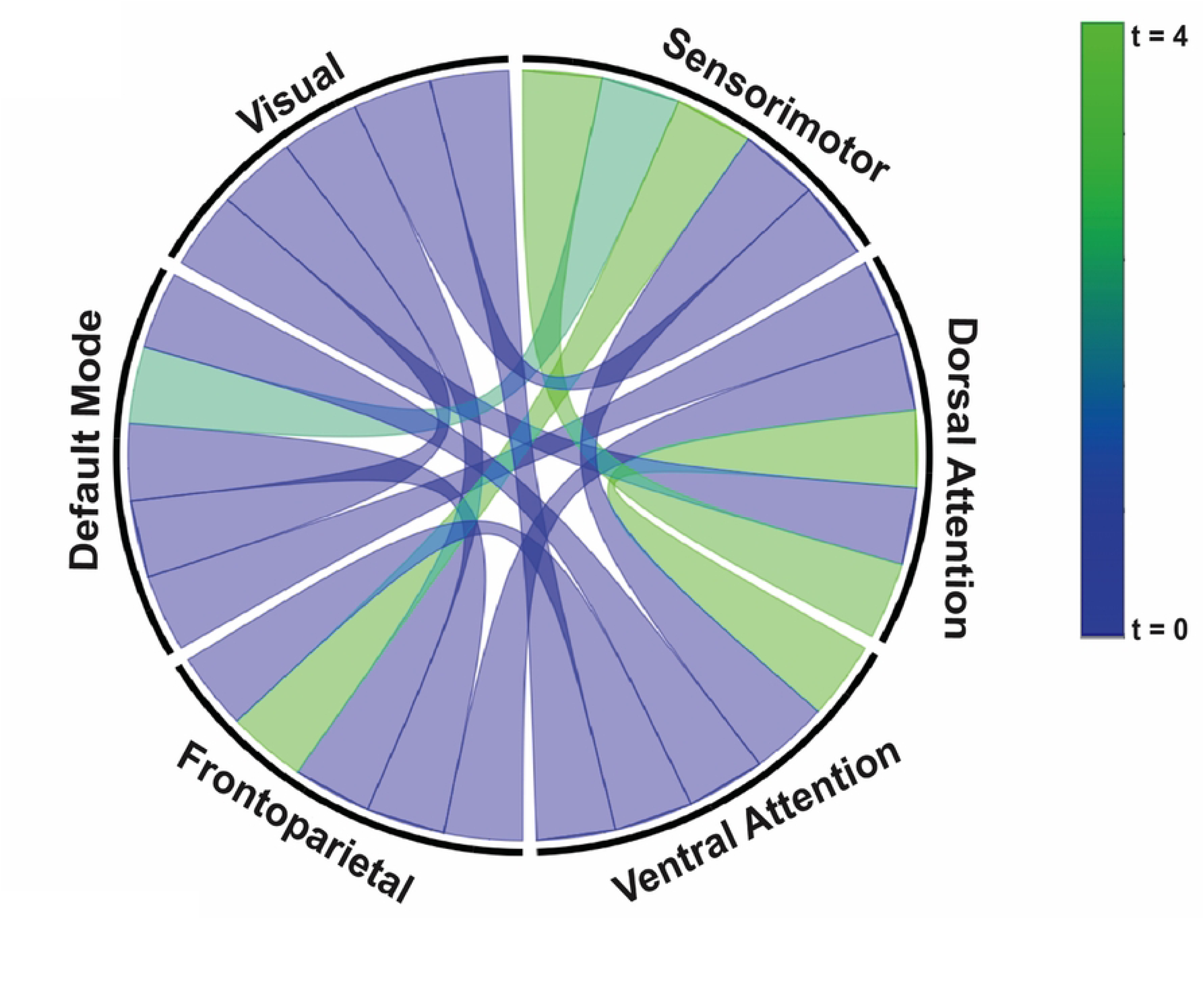
String plot showing significant post-training increases in resting-state functional connectivity between large-scale brain networks. *Note.* The color bar represents t-values, with warmer colors indicating stronger post-training increases.

## Discussion

Contemporary neuroscience increasingly views the brain as a complex, interconnected network rather than a collection of isolated modules. Specialized regions interact dynamically with one another, enabling seamless information processing and integration across structurally and functionally linked systems. Recognizing this complexity, emerging work has begun to move beyond isolated task-based activations to examine how training (or experience) reshapes the brain at the functional and structural network levels. In this research note, we examined neuroplastic changes in brain morphology, structural connectivity, and resting-state functional connectivity in response to a four-week EMST program in healthy young adults.

### Brain structural changes

Across both gray and white matter, no significant structural, morphometric (FWE-corrected *p* > .05) or connectivity-related (FDR-corrected *p* > .05) changes were observed after training, at least within the timescale and sample characteristics of the current study. While small decreases in total volume **(Table 2)** and whole-brain mean structural connectivity **(Fig 2A)** were observed from baseline to post-training, such fluctuations are common in longitudinal brain imaging and are often attributed to measurement variability, scanner noise, or preprocessing factors rather than true neuroanatomical change [23]. We believe that the following two factors may explain the absence of detectable widespread macrostructural changes. First, because EMST is a targeted intervention, its central neural engagement is likely restricted to specific neural circuits rather than widespread cortical or subcortical networks. Our previous work has demonstrated functional central adaptations [3,4], further supporting the idea that EMST activates select sensorimotor, motor learning, and cognitive pathways but does not necessarily drive broad, distributed engagement. Consistent with this finding, we also observed localized microstructural changes (increased fractional anisotropy) across several swallowing-implicated white-matter tracts, with the largest increase in the corticospinal tract [3]. These localized changes are noteworthy because similar tract-specific adaptations are commonly reported in the early stages of motor skill learning [24,25], suggesting that EMST, despite being categorized as strength training, may also engage skill-learning processes.

Second, the temporal dynamics of neuroplasticity of brain structure provide an additional explanation. Functional changes can emerge rapidly, but macrostructural and large-scale white-matter connectivity alterations typically require extended or more intensive training regimens [24,26,27]. While localized adaptations are characteristic of early stages of motor learning and training, longer or more intensive practice often leads to broader, distributed structural changes as the neural system consolidates and integrates the learned behavior. Furthermore, it is plausible that these structural changes are detectable in populations with greater neuroplastic potential, such as older adults or individuals with neurological impairment, warranting further investigation in these groups [28,29]. This possibility is particularly relevant given the limited sample size, pilot and exploratory nature of the current study, and potential ceiling effects in healthy young adults, all of which may have constrained our ability to observe macrostructural or large-scale white matter connectivity changes.

### Resting-state functional connectivity changes

In contrast to the stability observed in structural morphology and connectivity, the EMST program resulted in systematic alterations within select functional networks. First, we observed significant increases in whole-brain mean resting-state functional connectivity. Such increases are thought to emerge when specific, behaviorally relevant networks strengthen their internal and external coupling, thereby elevating global network statistics [30]. In the current study, the EMST-related enhancements likely reflect more efficient integration among select sensorimotor, cognitive, and attentional networks, thereby raising overall functional coherence. Such increases are consistent with models of early-stage neuroplasticity, in which targeted functional reorganization occurs early in training and precedes longer-term structural modifications [31].

Second, the graph-theory results provide converging evidence for EMST-induced reorganization of brain network topology. Trends toward increased global efficiency and clustering, together with significant increases in local efficiency and transitivity, suggest that the brain shifts toward a more integrated yet locally cohesive configuration following training. The observed reductions in modularity further reinforce this interpretation. Lower modularity reflects decreased segregation between functional communities, indicating that networks previously more distinct are becoming increasingly interconnected [30,32]. Collectively, decreases in modularity alongside increases in efficiency point to a transition toward a more globally integrated and flexible network architecture, consistent with early-stage functional reorganization preceding structural change.

Third, the EMST program strengthened connectivity both within and between select intrinsic functional networks, revealing more targeted forms of functional reorganization. Within-network increases in functional connectivity were most prominent in the sensorimotor, frontoparietal control, and dorsal attention networks, systems that showed increased functional activation in our previous work [3,4]. Strengthened connectivity within these networks may reflect enhanced sensorimotor processing, cognitive control, and attentional engagement, which are collectively recruited during EMST performance. These findings further support our earlier interpretation that EMST enhances cognitive–sensorimotor integration [4]. Between-network increases in functional connectivity reinforce this interpretation. Enhanced connectivity among sensorimotor, dorsal attention, frontoparietal, and default-mode networks indicates that EMST engages broader cross-system communication. Stronger sensorimotor–attention coupling suggests greater demands on sustained attention and performance monitoring, while increased dorsal–ventral attention connectivity reflects improved integration of externally directed attention with salience-driven control. Increased dorsal attention–frontoparietal coupling further points to heightened recruitment of multimodal circuits for attentional focus, sensorimotor feedback, and task regulation. Taken together, these findings indicate that targeted interventions such as EMST drive selective network reorganization that supports more efficient cognitive–sensorimotor integration, thereby offering mechanistic bases for developing precision therapies to remediate speech and swallowing deficits in ageing and disease.

### Conclusion and future directions

The current study aimed to investigate the effects of a four-week EMST program on structural and resting-state functional connectivity in healthy young adults using a multimodal imaging approach. Our findings demonstrate robust functional reorganization but no detectable macrostructural or large-scale white-matter connectivity changes following training. The absence of structural effects should not be interpreted as a lack of neuroplasticity. Functional plasticity often precedes macrostructural remodeling, and the current findings may represent the earliest stage of a longer sequence of neural change in which distributed structural or microstructural modifications emerge only with extended or more demanding training. Rather, it likely reflects the short duration of training, the targeted nature of EMST, and the limited structural malleability of healthy young adults. It remains plausible that structural changes would be more readily detectable in populations with greater neuroplastic potential. Future research could extend this work to healthy aging individuals and to clinical populations, including those with neurodegenerative diseases such as Parkinson’s disease, stroke survivors, and pediatric populations with developmental motor impairments.

The EMST program significantly increased resting-state functional connectivity, enhanced global and local network efficiency, reduced modularity, and strengthened within- and between-network coupling across sensorimotor, frontoparietal control, and dorsal attention systems. These changes closely parallel regions and circuits previously shown to exhibit EMST-related task-based activation, suggesting that early adaptations to EMST primarily involve functional reweighting and enhanced communication within and between cognitive–sensorimotor pathways. Given this pattern of strengthened within- and between-network coupling, future work would benefit from effective connectivity approaches, such as dynamic causal modeling, to clarify the directional flow of information. Such analyses could confirm whether EMST primarily enhances top-down influences from cognitive-control networks, bottom-up sensorimotor drive, or a bidirectional heterarchical network that supports improvements in upper aerodigestive functions.

## Data Availability

The data that support the findings of this study are available from the corresponding author upon reasonable request.

## Acknowledgments

This work is supported by the Council of Academic Programs in Communication Sciences and Disorders (CAPCSD) scholarship to RK.

## Supporting information

**S1 Table.** Means, standard deviations and results of the paired t-test comparing surface-based morphometric measures for the study’s region of interest (ROIs) across baseline and post-training conditions.

**S2 Table.** Mean, standard deviation and paired t-test results comparing nodal-level structural connectivity network measures for specific ROIs across baseline and post-training conditions.

**S3 Table.** Mean, standard deviation and paired t-test results comparing nodal-level functional connectivity network measures for specific ROIs across baseline and post-training conditions.

